# Brain-to-brain coupling forecasts future joint action outcomes

**DOI:** 10.1101/2023.09.21.558775

**Authors:** Roksana Markiewicz, Katrien Segaert, Ali Mazaheri

## Abstract

In this study, we investigated whether brain-to-brain coupling patterns could predict performance in a time-estimation task which requires two players to cooperate. The participant pairs, were tasked with synchronising button presses after converging on a shared representation of ’short’, ’medium’, and ’long’ time intervals while utilizing feedback to adjust responses. We employed EEG-hyperscanning and focused on post-feedback brain activity. We found that negative feedback led to increased frontal mid-line theta activity across individuals. Moreover, a correlation in post-feedback theta power between players forecasted failed joint action, while an anti-correlation forecasted success. These findings suggest that temporally coupled feedback related brain activity between two individuals serves as an indicator of redundancy in adjustment of a common goal representation. Additionally, the anti-correlation of this activity reflects cognitive strategic mechanisms that ensure optimal joint action outcomes. Rather than a *paired overcompensation*, successful cooperation requires *flexible strategic agility* from both partners.

## Introduction

Successful joint action entails sharing a common goal and striving towards it, while also incorporating feedback when deviations from the goal occur. To successfully cooperate, it is thus crucial to process feedback and adjust behaviour accordingly. While EEG-hyperscanning has previously been used to investigate joint action (e.g., Hamilton, 2021), no study to date has investigated the brain-to-brain coupling of neurocognitive mechanisms that drive the adjustment of shared representations of the common goal in response to feedback. In the current study, we investigate the specific brain-to-brain activity, in response to negative feedback, that can forecast the strategic cognitive mechanisms underlying adjustment of the shared representation of the task and facilitate successful performance in joint action cooperation.

The primate’s brain has evolved to not only manage the demands related to cognitive abilities but more importantly to navigate and work with large social groups (Dunbar, 1992). Joint action is a type of social interaction, in which a shared goal is achieved through the coordinated actions of at least two individuals in time and space (Sebanz et al., 2006). Joint action is present in everyday life, for example when moving a heavy sofa up a flight of stairs. The success of joint action performance is reliant on creating shared representations, the ability to predict actions as well as the outcomes of one’s own and others’ actions, including adjustments made when the outcome of an attempt is unsuccessful (Sebanz et al., 2006). A first mechanism that underlies joint action is about ensuring that the coordinating partners can *guide attention* to perceive the same event or object (like creating common ground).

Secondly, for successful joint action to occur, *action observation* is required. A corresponding representation of the object/event of interest in the observer’s action system is created. This aids the understanding of the action, creates ‘common ground’ in terms of action goals, and supports prediction of each other’s action outcomes. For example, (Flanagan & Johansson, 2003) observed that the gaze of the action observer precedes the action of their partner. Lastly, but perhaps the most important, is *action adjustment,* which refers to adjusting behaviour in space and time to complement the action of the coordinating partner. For example, lowering your end of the sofa when you see your moving partner is getting stuck with their end (i.e., the results of the adjusted shared representation of the common goal). Joint action is influenced not just by an individual’s beliefs about their own abilities but also by the beliefs about what they can achieve in collaboration with others (Marsh et al., 2006). In the present study, we will focus on using feedback to make the adjustments needed to increase joint action success. To do this, we will examine the dynamic relationship between interacting brains (Dumas, 2011; Hari et al., 2015; Konvalinka & Roepstorff, 2012; Stolk et al., 2016) using a hyperscanning approach.

In the last decade, the social neuroscience field has seen a trend towards using a ‘multi brain framework’ to study how humans interact with each other. Hyperscanning involves the simultaneous recording of brain activity (either hemodynamic or neuroelectric) in multiple individuals during interactive task performance. It allows for investigations on how the activity dynamics of two brains (or more) underlies the continuous adaptation of one’s actions in response to the changes in actions of someone else. Since first implemented by Montague et al. (2002), two-brain science has been used to study classroom dynamics (Dikker et al., 2017), communication (Schoot et al., 2016; Stolk et al., 2013), music (Babiloni et al., 2012), and coordinated button presses (Balconi & Vanutelli, 2017; Cui et al., 2012; Funane et al., 2011; see Czeszumski et al., 2020; Konvalinka & Roepstorff, 2012 for detailed reviews). Also, interpersonal dynamics such as cooperation or joint action have widely been studied using hyperscanning methods (see Balconi & Vanutelli, 2017). Joint action relies on the postulation of anticipating and predicting other people’s actions and adjusting our own actions accordingly (Decety & Sommerville, 2003). ‘Coupling’ between individuals can occur on motor, perceptual or cognitive levels during joint action (Knoblich et al., 2011) with behavioural synchrony often reflected in neural synchrony. This neural connectivity as measured by hyperscanning is the correlation between neurophysiological signals that are temporally related but spatially distinct (i.e., in two separate brains) (Dumas et al., 2012).

EEG hyperscanning is particularly useful in the context of the present study due to its high temporal resolution, which allows for a real-time window into the neural processes induced by feedback in the unfolding dynamics of social interaction on a millisecond scale. Several EEG hyperscanning studies have reported inter-brain synchronous oscillatory patterns in joint action contexts, both in terms of oscillatory power as well as phase during fine motor coordination (Dumas et al., 2010; Tognoli et al., 2007), verbal coordination (Ahn et al., 2018) and other complex joint actions (Astolfi et al., 2012). The mechanisms that support adjustment of the shared representation of the task goal in response to feedback in a joint-action context are key to explaining joint action outcomes, but they are not clear to us yet.

Only two studies to our knowledge have investigated the inter-brain dynamics in tasks with an incorporated feedback element. Mu et al. (2016) reported on the heightened inter-brain alpha phase synchrony *during* a coordination task (vs control), but not in response to feedback. Conversely, Balconi et al. (2018) examined directly the effect of external feedback on joint action in the context of EEG inter-brain dynamics. However, in this study, feedback was superficially created and not based on the actual joint action performance. From this, it is evident that there is a clear gap in the literature: the precise neural mechanisms that facilitate adaptive responses to feedback and thus shared representation of the common goal within the context of joint action remain elusive.

The impact of feedback has been studied extensively in tasks that do not involve a joint-action component (e.g., van de Vijver et al., 2011), or, in tasks that involved a joint-action component but only examined the neural dynamics of one of the participants (e.g., Czeszumski et al., 2019; Itagaki & Katayama, 2008; Picton et al., 2012). A careful look at these studies enables us to propose precise predictions about the inter-brain mechanisms that might be involved in post-feedback adjustments in a joint-action setting. It has been established that a feedback cue induces oscillatory EEG changes, time-locked, but not necessarily phase-locked, in the 3-7 Hz (theta range) with a maximal midline frontal distribution. This feedback induced neural signature is thought to reflect the initiation of response adjustments (i.e. cognitive control) overriding ‘status quo’ responses (Cavanagh et al., 2013; Cavanagh & Frank, 2014). The midline theta response can be reliably measured on a trial-by-trial basis, providing valuable insights into the neural-dynamics subserving behavioural adjustment. This measure affords a real-time window into how the brain adapts and adjusts behaviour based on feedback (van de Vijver et al., 2011), allowing us to explore the intricate relationship between neural activity and behavioural responses. In the present investigation, our focus lies on theta activity as a reflection of adjustment in the mental representation of the time intervals between the partners within a pair. However, rather than solely examining how the amplitude of feedback related theta activity in each participant (referred to as "player") relates to their performance in future trials, we, by using EEG hyperscanning, are specifically interested in how the *correlation* between these responses can effectively forecast their joint action success.

We implemented an EEG hyperscanning approach while participants completed an innovative time-estimation task. The objective of this task was to converge on a mutual representation of what a ‘short’, ‘medium’ and ‘long’ interval entailed. Specifically, auditory cues instructed participants to either press a button after what they mutually judged to be a short, medium or long time. Participants reached a successful joint action outcome if they reached a shared representation of the time intervals and thus converged on their responses.

Importantly, feedback was given after each trial with respect to their joint action outcome, and if unsuccessful, about who was faster and by how much. With this, we created an ideal situation to study the inter-brain mechanisms that support feedback processing and subsequent behavioural adjustment. By utilizing this time-estimation task, we were able to study how individuals develop shared understanding and mutually converge on an idea (in this case of timing intervals) exclusively through the external feedback they received. This task included well-defined joint action outcomes, namely successful and failed cooperation, enabling us to investigate the separate neural mechanisms underlying these distinct outcomes in joint action.

In our theoretical framework, we hypothesize that the absence of strategy alteration or mutual adjustment in the mental representation of the time intervals (short, medium, and long) in both players, we refer to this as *strategic stagnation*, would result in unsuccessful joint action. Conversely, if both partners actively modify their strategies and adapt their mental representations of the time intervals, this could potentially facilitate adaptive adjustments and promote successful joint action.

However, the risk of both players attempting to adjust their strategies is that there would be a redundancy of adjustment in the mental representations of the time intervals, which could lead to *paired overcompensation* and thus maladaptive behavioural adjustment. Therefore, our alternative hypothesis posits that joint action success is more likely when players exhibit *flexible strategic agility*. In this scenario, one partner adjusts their mental representation to a greater extent than the other, allowing for asymmetry in their adaptation process. This asymmetry may manifest as one player making more pronounced changes in their mental representation compared to the other, leading to effective coordination and ultimately contributing to successful joint action. We theorise that this mechanism may strike an optimal balance, characterized by flexible strategic agility and asymmetrical adjustments in the mental representation of time intervals, and thus enable a balanced mutual convergence that facilitates successful joint action.

## Results

Cooperation was measured via a non-verbal task, completed in pairs, while EEG was recorded from both players simultaneously (see Figure 1 for experimental set up and trial presentation). Participants synchronised their button presses with their partner within the pair. Auditory stimuli (high, medium, or low pitch tone) at the start of each trial indicated the duration of time (referred to as ‘short’, ‘medium’, or ‘long’) that participants had to wait before pressing their button and attempting to synchronise it with the other player. Participants adjusted their representations of short/medium/long and thus their button-press responses solely based on the feedback provided at the end of each trial.

**Figure 1.**
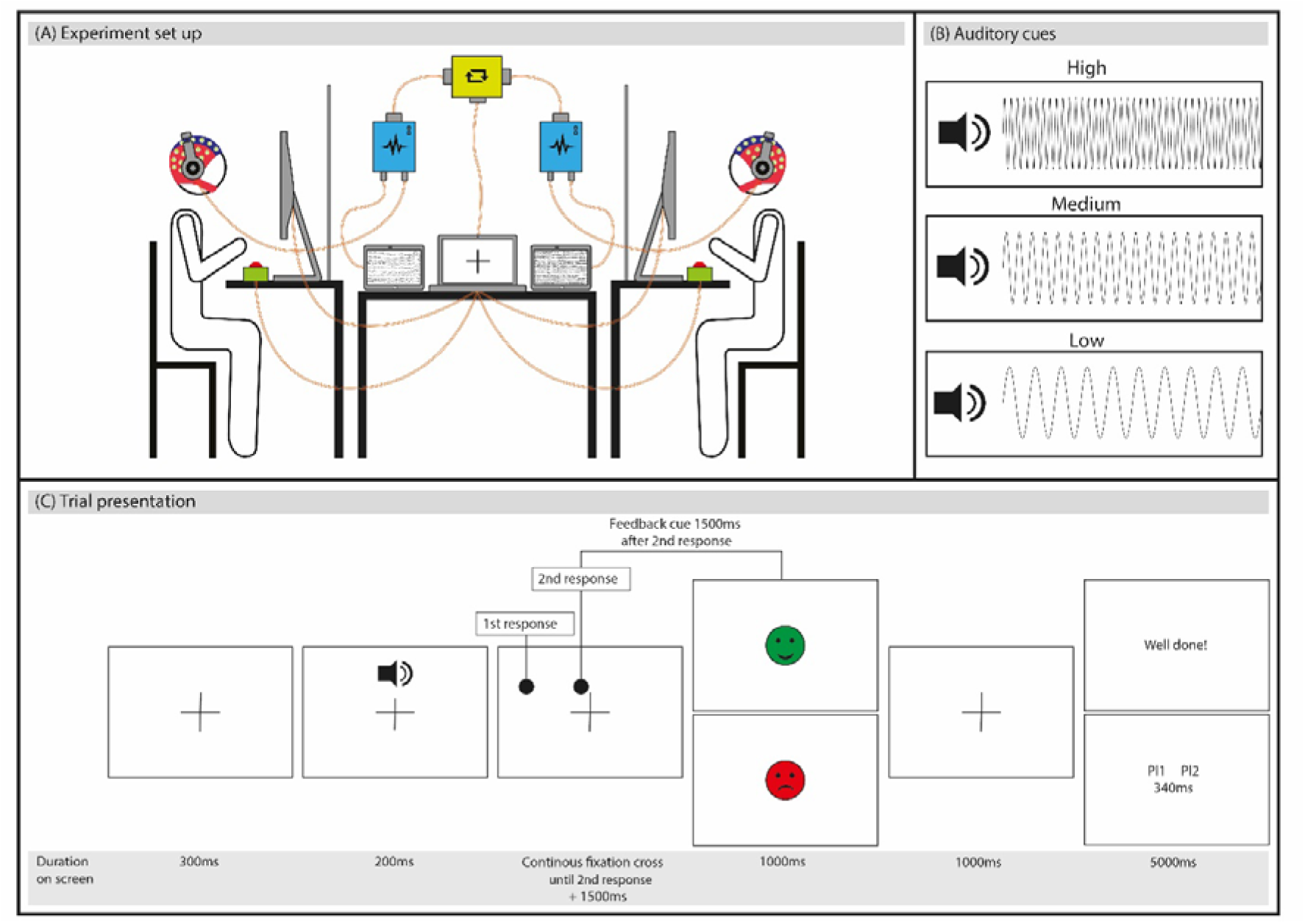
The experimental setup and trial presentation. (A) Participants wore Waveguard caps with 32 cap-mounted Ag/AgCI electrodes. The ANTneuro EEGsports amplifier systems amplified the signals for each participant. The EEG signals were synchronized using a parallel port splitter. The stimuli were presented on a central laptop connected to two identical monitors. Participants listened to auditory cues through noise-cancelling headphones and used a button box to record their responses. (B) The experiment used three auditory cues: high, medium, and low frequency tones, corresponding to short, medium, and long waiting times. (C) Trial presentation involved both participants receiving the same stimuli simultaneously. The order of the button responses was completely driven by the participants and thus resulting in trial-to-trial variations of who pressed first/ second. The feedback stimuli varied based on trial outcome: a green smiley face and "Well done" for successful cooperation trials, and a red sad face with feedback text indicating which player pressed the button first/second and the elapsed time for failed cooperation trials. Participants were informed of their player number to understand the feedback text.

### Participants adhered to task instructions

We used R (R Core Team, 2021) and *lme4* (Bates et al., 2015) to perform a linear mixed effects analysis of the relationship between condition type (short, medium, and long) and response times of the individuals. This was to check whether participants adhered to task instructions of waiting short, medium, and long amounts of time before synchronising their button presses (depending on the auditory cue type). We included a fixed effect of condition type (short, medium, and long), and random effects of intercepts for pairs and subjects, as well as by-pair and by-subject random slopes for the effect of the condition type. When a model did not converge with the maximal random effects structure, we simplified the random slopes until convergence was reached. The final and simplified model specification was as follows: RT ∼ Condition + (1|Pair) + (1|Subject). Table 1 reports the model output. We used custom contrasts and thus compared conditions *short* vs. *medium* and conditions *medium* vs. *long* in one model (model A), and conditions *short* vs. *long* in another model (model B). Figure 2 shows the average response times (sec) and accuracy (%) for each condition across all participants, Suppl. Fig. 1 displays the response times of player 1 and player 2 of a representative pair in function of trial number for each condition.

**Figure 2.**
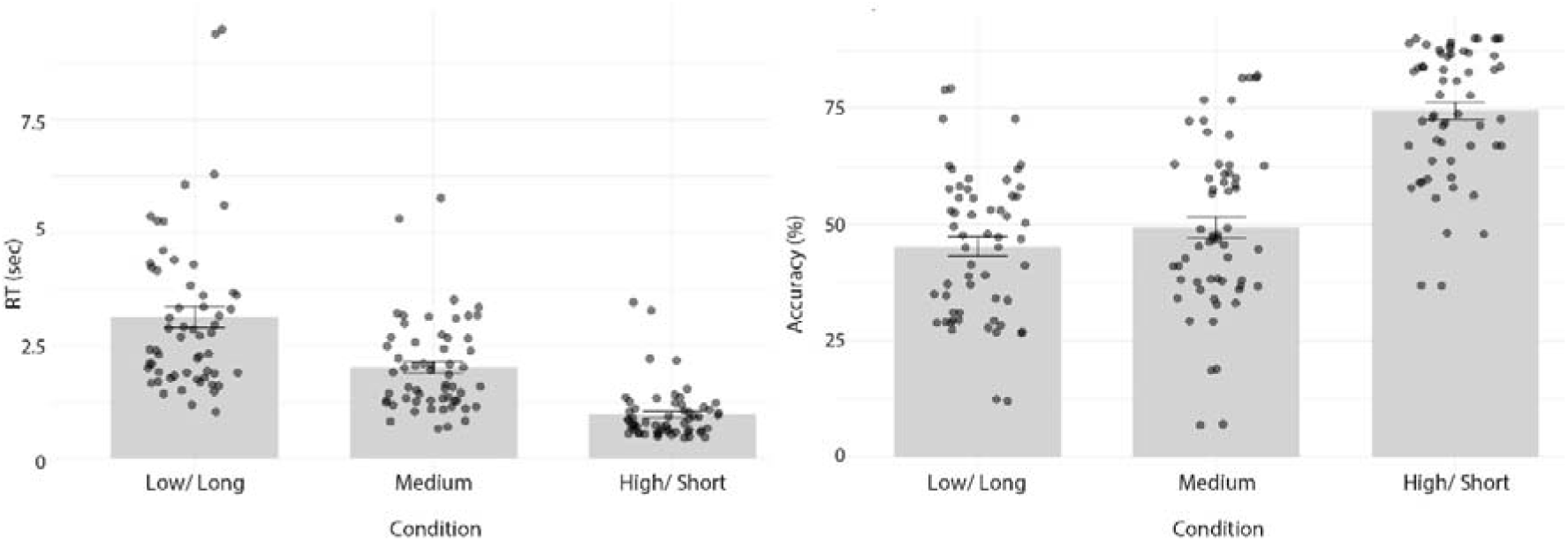
Average response times in seconds (left), and average accuracy in percentages (right) per condition across all participants. The error bars represent standard error and the black dots represent individual data points.

**Table 1.** Summary of the mixed effects models for the Reaction Times.

**Model A**
|  | Estimate | SE | *t* value | *p* value |
| --- | --- | --- | --- | --- |
| Intercept | 2.077 | 2.039 | 10.19 | <.0001 |
| Condition *short* vs Condition *medium* | -1.106 | 1.524 | -72.58 | <.0001 |
| Condition *medium* vs Condition *long* | 1.049 | 1.524 | 68.83 | <.0001 |

**Model B**
|  | Estimate | SE | *t* value | *p* value |
| --- | --- | --- | --- | --- |
| Intercept | 4.176 | 2.039 | 20.48 | <.0001 |
| Condition *short* vs Condition *long* | -2.15 | 1.524 | -141.42 | <.0001 |

N = 17398

Additionally, we examined whether the outcome of each trial (i.e., the time difference between the button presses) could be predicted based on the interplay of behavioural adjustments made by both players after the previous trial. To do this, we computed a multiple linear regression model, with the predictor variables being: the behavioural adjustment of player one and behavioural adjustment of player two, as well as their interaction. The outcome variable was the cooperative outcome (i.e., the time difference between the button press of player one and two); cooperative outcome ∼ behavioural adjustment of player one * behavioural adjustment of player two. We found that the model explained a statistically significant but weak proportion of variance (F(3, 2647) = 61.40, p < .001), and importantly the interaction of behavioural adjustment of player one and player two was a significant predictor of cooperative outcomes (β = -0.16, p < .001). This suggests that the cooperative outcome is shaped by the dynamic interaction of behavioural adjustments made by both players (rather than solely their individual adjustments). See the supplementary material (section: *Interplay of behavioural adjustment (between players) was predictive of cooperative outcome)* for how behavioural adjustment was calculated and more detailed results).

### Feedback related oscillatory modulations in the theta band

We compared the TFRs of power between successful and unsuccessful cooperation conditions across all participants (N=59) aligned to the onset of the feedback text. Figure 3 displays the TFRs of power locked to the onset of the feedback text for the incorrect and correct conditions, and the topographic distributions of the condition effect in the theta band.

**Figure 3.**
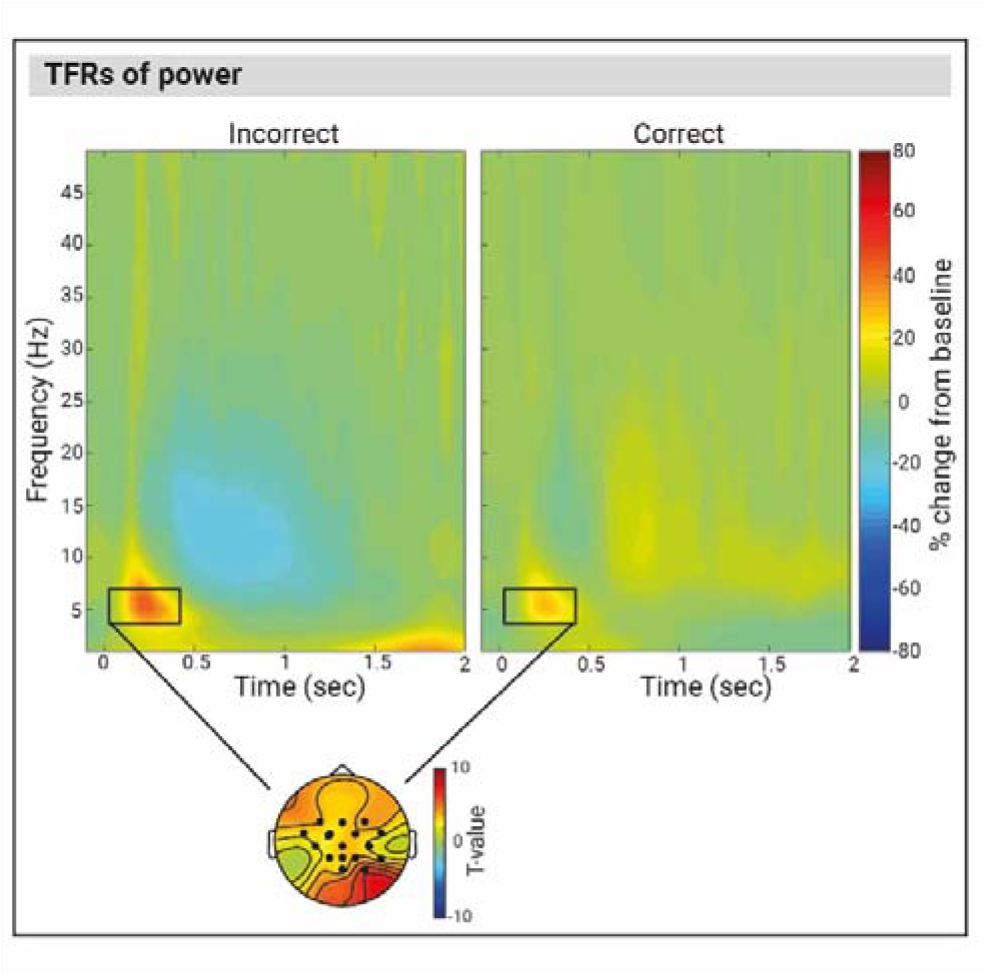
Time Frequency representations of power at the Cz electrode for unsuccessful trials (left), and successful trials (right) averaged over all individuals (N=59), locked to the onset of the feedback text. The topoplot illustrates the clusters of electrodes that show the most pronounced condition difference. Black rectangles indicate a significant condition difference (p < .05, cluster corrected) in the theta band.

Below we describe the significant oscillatory differences between successful cooperation and unsuccessful cooperation conditions that were identified using non-parametric cluster-based permutation in the theta band (4-7Hz) post-feedback text onset.

We observed significant condition differences (p = .003) in post-feedback theta activity between the successful cooperation and unsuccessful cooperation condition, which was maximal over the central and centro-parietal channels. Specifically, theta power was stronger right after the onset of the feedback text in the failed cooperation condition until around 0.4sec compared to the successful cooperation condition.

Further, we also observed significant condition differences in the feedback related activity in the delta (1-4Hz), alpha (8-14Hz), low beta (15-20Hz) and high beta (20-25Hz) frequency bands as well as ERP differences (see supplementary materials and Suppl. Fig. 1 for a comprehensive description of all effects).

### Inter-player theta power coupling forecasts cooperative outcome on the next consecutive trial

We compared the inter-player power coupling (see section *Methods: Forecasting cooperative outcome analysis* for how the inter-player power coupling was calculated) between two conditions: *forecasting unsuccessful cooperation* (i.e., failed cooperation trials followed by *a consecutive* unsuccessful cooperation trial) *and forecasting successful cooperation* (i.e., failed cooperation trials followed by *a consecutive* successful cooperation trial) using non-parametric cluster-based permutation tests across all pairs (N = 29 pairs). Figure 4 displays inter-player power coupling for (A) *forecasting unsuccessful cooperation* and (B) *forecasting successful cooperation*. We found an inter-player oscillatory power coupling signature that forecasts cooperative outcome on the next consecutive trial (i.e., local effect). Using the non-parametric cluster-based permutation tests (with a pre-defined time window of 0 - .4sec post-feedback based on the feedback related results in the Time Frequency domain), we observed a significant condition effect (p = .006) in the theta frequency band (this effect was also observed when an exploratory time window of 0 to 2sec was used in the non-parametric cluster-based permutation tests, p = .051). The effect corresponded to a cluster that spanned between .15 to .2sec and was maximal over two electrodes (Fz and FC2) that neighboured each other. The inter-player theta power coupling showed the opposite patterns of results in the two conditions. Forecasted cooperative failure was associated with a positive correlation between theta power of player one and player two (i.e. the greater the theta power in one player, the greater the theta power in the other player within a pair, or, the smaller the theta power in one player, the smaller the theta power in the other player). On the other hand, forecasted cooperative success was linked with a negative correlation between the theta power of player one and player two (i.e., the greater the theta power of one player, the lower the theta power of the other player within a pair or vice versa).

**Figure 4.**
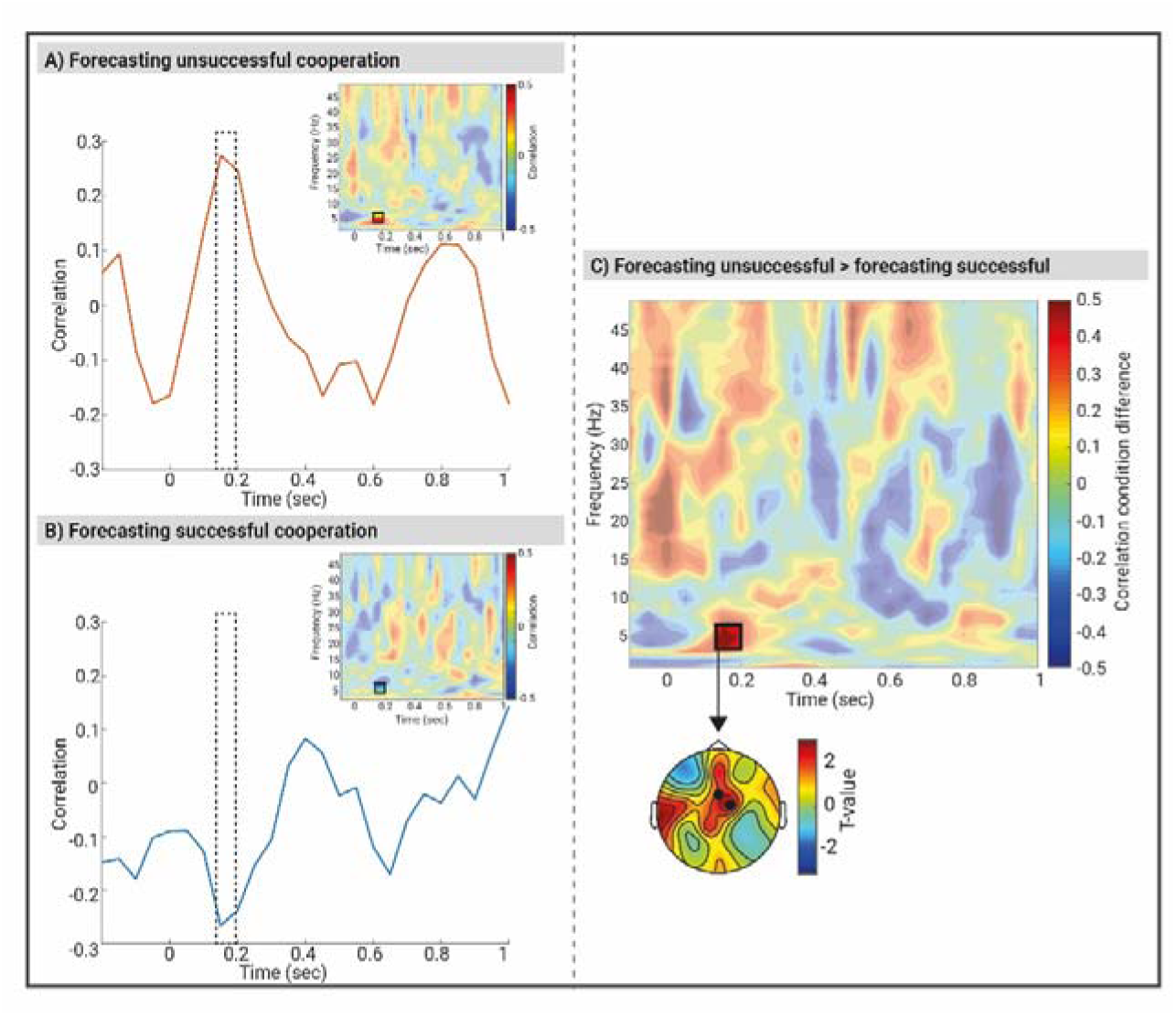
Forecasting joint action outcomes (successful vs. unsuccessful cooperation) on next consecutive trials based on the correlation values (Fisher Z) of oscillatory power coupling between player 1 and player 2 after the onset of feedback text in the failed cooperation condition. Line plots are representing the correlation (Fisher Z) values averaged over Fz and FC2 electrodes in the theta frequency band (4-7Hz) forecasting (A) unsuccessful cooperation trials and (B) successful cooperation trials. The dashed rectangles indicate the time windows of the significant between condition differences (as calculated via the non-parametric cluster-based permutation tests). The heatmaps are displaying the correlation values (Fisher Z) of oscillatory power coupling averaged over Fz and FC2 electrodes between player 1 and player 2 locked to the onset of the feedback text forecasting (A) unsuccessful and (B) successful cooperation. The black rectangles indicate significant condition differences (p < .05, cluster corrected). (C) A heatmap is representing the condition difference (forecasted unsuccessful cooperation – forecasted successful cooperation) correlation values (Fisher Z) of oscillatory power coupling (averaged across the electrode clusters that indicate maximal condition difference, i.e., Fz and FC2) between player 1 and player 2 locked to the onset of feedback text. The black rectangles indicate significant condition differences (p < .0.05, cluster corrected).

### Inter-player coupling in delta, alpha, or beta bands does not forecast cooperative outcome on the next consecutive trial

We did not find any significant inter-player power coupling signatures of forecasting cooperative outcomes in the alpha band using the pre-defined .3 to 1.75sec time window. Further analyses using exploratory time window of 0 to 2sec post-feedback text also did not yield significant effects in any of the frequency bands (delta, alpha, low beta, and high beta).

### Relationship between behaviour and theta dynamics on a trial-by-trial basis

The above analysis (i.e., *Inter-player theta power coupling forecasts cooperative outcome on the next consecutive trial)* treated cooperative outcomes as dichotomous variables defined by a pre-determined threshold: cooperative failure vs. success. However, we also sought to examine whether the trial-by-trial inter-player theta dynamics are predictive of *continuous* cooperative outcomes on behavioural level (i.e., the absolute difference in RT’s between player one and player two). We carried out a multiple linear regression analysis as follows: cooperative outcome RT difference ∼ averaged theta power of player one * averaged theta power of player two + absolute theta power difference between players (see *Trial-by-trial analysis linking theta dynamics with behavioural outcomes* in Methods for further details). The model explained a statistically significant and weak proportion of variance (F(4, 2738) = 7.64, p < .001). The model revealed that the averaged theta power of player one (β = 0.08, p < .001), and player two (β = -0.02, p = 0.008) were significant predictors of cooperative outcome. Importantly, the model showed that the interaction of averaged theta power of player one and player two (i.e., averaged theta power of player one * averaged theta power of player two) significantly predicted the continuous cooperative outcome (β = -0.25, p < .001). This suggests that cooperative outcomes are significantly driven by the dynamic interplay of theta power modulations of *both* players.

### Feedback related modulations in theta band do not reflect the required type of individual adjustment

We aimed to further explore the underlying mechanism responsible for the observed brain-to-brain activity in the theta power forecasting joint action outcome. Here, our aim was to examine whether the observed inter-player coupling effect originates from the *mutual* adjustment or whether it is attributed to a singular process performed independently by each individual (and thus the brain-to-brain coupling effect is merely a by-product of cognitive processing). To achieve this, we compared trials based on the individual type of adjustment required for the following trial; *unsuccessful cooperation trials that necessitated speeding up* in the subsequent trial vs *unsuccessful cooperation trials that necessitated slowing down* in the subsequent trial.

The non-parametric cluster-based permutation tests did not yield any significant condition differences in the theta power between the unsuccessful trials that required speeding up vs. slowing down. In other words, at the *individual* level, theta power did not serve as a predictive factor for determining the type (speed up or slow down) of behavioural adjustment required in the subsequent trial. Thus, we can be confident that the brain-to-brain effects reported in the *Inter-player theta power coupling forecasts cooperative outcome on the next consecutive trial* section arise from the mutual adjustment of time interval representation in this joint action context and not merely as a by-product of common cognitive processing.

## Discussion

In this study, we investigated whether brain-to-brain coupling patterns in a two-player game could predict performance in a joint-cooperation task. The aim of the task for the players was, upon hearing auditory cues, to establish a shared understanding of ‘short’, ‘medium’, and ‘long’ intervals and synchronise their button presses with each other. The outcome of the joint action was successful only when players converged on their responses. We found that receiving negative feedback led to increased frontal mid-line theta activity across individuals. Moreover, a correlation in post-feedback theta power between players forecasted failed joint action, while an anti-correlation forecasted success. We suggest that the temporally coupled feedback related brain activity between two individuals is an indicator of one of two strategies, either redundancy in adjustment of their representation of the task goals (i.e., *paired overcompensation;* mutual high theta power) or lack of adjustment from both partners (i.e., *strategic stagnation*; mutual low theta power). On the other hand, the anti-correlation of this activity reflects cognitive strategic mechanisms that ensure optimal joint action outcomes. We propose that our results indicate that successful joint action outcomes are achieved by the implementation of *flexible strategic agility*, whereby in a given trial one partner within the pair demonstrates greater adaptability than the other (the higher the theta power in one player, the lower the theta power in the other player). This flexible adaptive approach emerges as a key determinant for converging on the representation of the task goals and thus fostering successful cooperation.

Our findings demonstrate how the dynamics of the brain activity induced by feedback between two players could forecast joint action outcomes. With this, we filled a crucial gap in the literature by emphasizing the significance of interpersonal neural patterns in predicting joint action outcomes on subsequent trials by focusing on the correlation of the brain-to-brain neural responses, rather than examining the feedback related neural responses on an *individual* level.

The use of the external feedback allowed the players to adjust their representations of the timing intervals and therefore responses for future trials. On an *individual* level we found distinct patterns of neural activity related to feedback: an increase in frontal midline theta activity after negative feedback and an increase in centro-posterior midline beta following positive feedback (see supplementary material for more details on feedback related modulations in *delta, theta, alpha, and beta* bands). The enhanced theta following negative feedback is linked with enhanced cognitive control and the need for behavioural adjustment following an error (Cohen, 2016; Cohen et al., 2007; Spitzer & Haegens, 2017), whereas the increased beta following positive feedback signals the requirement for strategy maintenance after a correct response. Furthermore, our analysis revealed that at the *individual* level, no significant differences emerged in theta power modulations between unsuccessful trials that required speeding up compared to those that required slowing down in the subsequent trial.

Most importantly, we found the correlation of brain-to-brain activity in the feedback related theta power to be a predictor of cooperative outcomes on subsequent trials. Given the well-established role of theta oscillations in adjustment following negative feedback, wherein they override ‘status quo’ responses at individual level, it is reasonable to anticipate distinct brain-to-brain patterns within the theta frequency range, that are predictive of future joint action outcomes at a paired level. Specifically, *correlated* brain-to-brain activity in the theta power significantly increased the likelihood of an error in the subsequent trial. We speculate that this can be attributed to two potential unsuccessful cognitive strategies adopted by the players. We suggest a first unsuccessful strategy is for the two players to passively apply *strategic stagnation*, where neither of the players adjusted their representation of the timing intervals, resulting in failed joint action. Interestingly, an alternative yet equally unsuccessful strategy, which we suggest is a concerted effort of strategy modification from both players (i.e., *paired overcompensation)*, resulted in the redundancy of adjustment of the representation of the task goals, overcorrection, and thus subsequent failed joint action. A strategy towards success on the other hand, was *anti-correlated* brain-to-brain activity in the theta power, which forecasted successful subsequent trials. Here, our proposed interpretation is that players employed a *flexible strategic agility,* wherein one partner adjusted their cognitive strategy and the representation of the common goal to a greater extent than the other (see Figure 5 for a schematic representation of the results). This *flexible strategic agility* facilitated a harmonious balance of adaptive adjustments, resulting in mutual convergence of the time intervals, and thus successful cooperative outcome in the subsequent trial.

**Figure 5.**
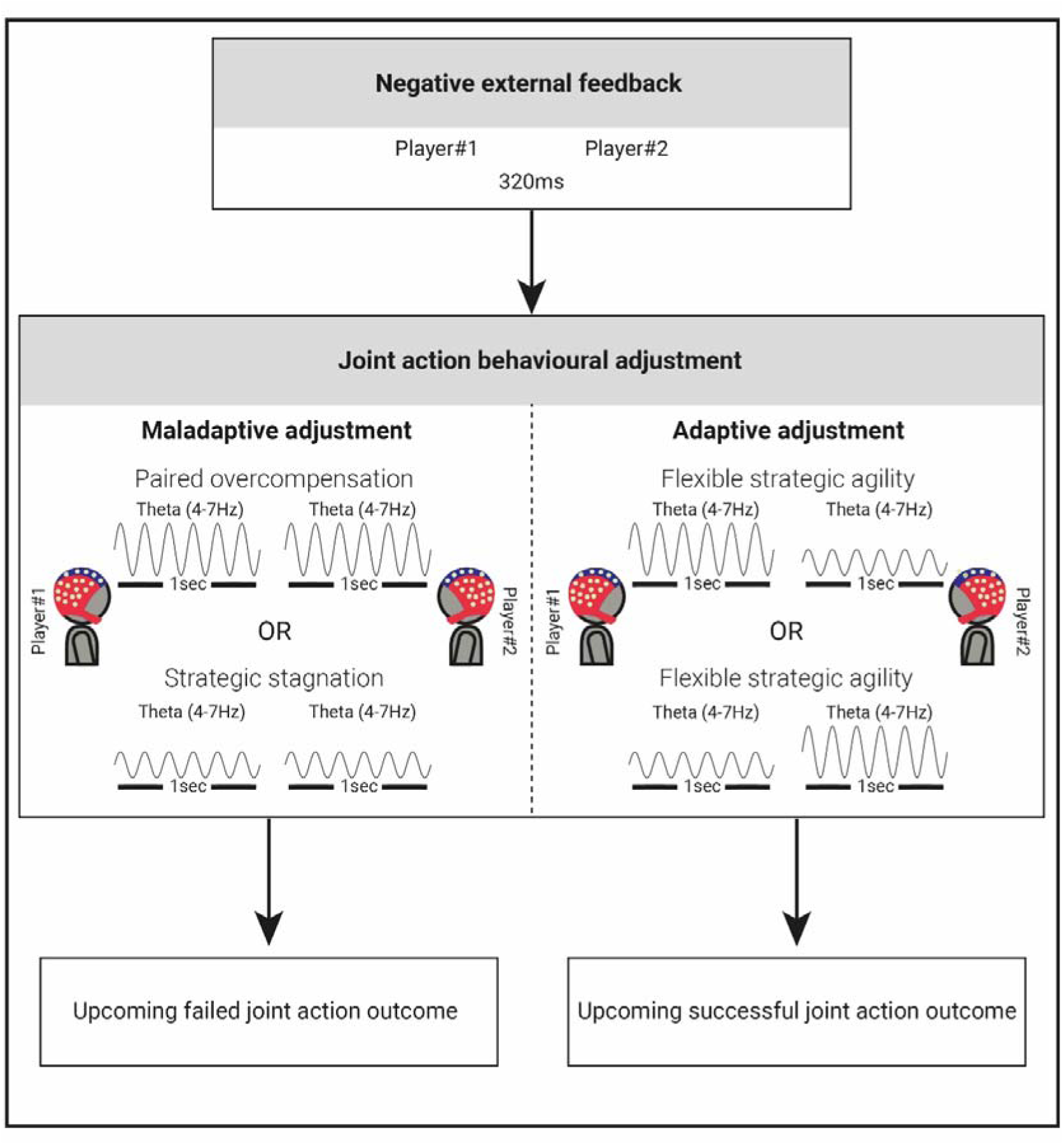
Schematic representation of the theta power modulation dynamics between individuals following feedback.

Our findings are consistent with the traditional view of enhanced theta power being linked with enhanced cognitive control (Cavanagh et al., 2009, 2010; Cohen, 2011; van de Vijver et al., 2011). Specifically mid-frontal theta is thought to be a neural marker of the adaptive response to (potential) errors (Cavanagh & Frank, 2014; Cohen, 2014, 2016). Oscillatory activity in the theta band in the frontal network (including the medial frontal cortex (MFC) and lateral prefrontal cortex (pFC)) is linked to the monitoring of ongoing actions and signalling unfavourable action outcomes. A greater theta power increase has consistently been observed following errors compared to successful outcomes (Cavanagh et al., 2009, 2010; Luu et al., 2004). This theta band activity also extends its role to predicting learning from negative feedback suggesting its involvement in behavioural adjustment in response to feedback that arises from suboptimal actions and thus outcomes (van de Vijver et al., 2011). Specifically, the mPFC theta increase has been linked to predicting post-error reaction time slowing, suggesting greater engagement of cognitive control for identifying and fixing errors (Crivelli-Decker et al., 2018). Conversely, beta oscillatory power increases have been observed following correct responses and positive feedback (Cohen et al., 2007; Marco-Pallares et al., 2008, 2009) indicating the need for strategy maintenance (Engel & Fries, 2010; Spitzer & Haegens, 2017). This beta band power change facilitates the reinforcement of the current motor or cognitive strategy via top-down influence (van de Vijver et al., 2011). We replicated and further extended literature findings in this field by demonstrating that modulations in theta and beta band activity in response to feedback are also elicited in joint action settings with the task goal being mutual convergence of timing intervals, rather than individual learned responses.

In addition, our inter-brain findings expanded the traditional view on the link between theta power modulation and cognitive control and moved this framework beyond the potential predictive nature of oscillatory activity on the individual level. Our findings demonstrated that we could apply the current theory to a joint action scenario where its success is reliant upon the convergence and creation of a mutual representation of time intervals between individuals based on the feedback provided. This view is further supported when switching from the dichotomous nature of the joint action outcome (i.e., successful vs. unsuccessful cooperation), and instead examining joint action outcomes as an absolute time difference between players on a trial-by-trial basis. We demonstrated that the interplay of the averaged theta power of player one and player two was a significant predictor of cooperative outcome as measured as a continuous variable (i.e., absolute difference in RT’s between the players). The trial-by trial brain dynamics of two interacting players and their subsequent behavioural performance suggests a mechanistic role of the brain signatures we observed for cooperative outcomes.

We revealed distinct strategies that accompanied different joint action outcomes: successful joint action and failed joint action. We demonstrated that flexible joint action adjustment, forecasted future successful joint action. We propose that employing a *flexible strategic agility* allowed for the mutual convergence of shared representations of timing intervals to be developed amongst the players. The *flexible strategic agility* emerged when varying levels of adaptability among partners were developed, with one player exhibiting a greater propensity for strategic adjustment than the other. This could then lead to the harmonious balance of adaptive adjustment, and thus successful cooperation. We posit that this is displayed in the anti-correlated brain-to-brain activity, wherein the higher the theta power in one player, the lower the theta power in the other player. We speculate that this *anti-correlated* brain-to-brain activity was indicative of the players engaging in rather complementary adjustment processes and increasing the likelihood of uncovering different solutions for convergence on a representation of the common goal.

On the other hand, a correlated brain-to-brain activity in the theta power was foretelling of future failed joint action. We speculate that this could be attributed to the adoption of two distinct cognitive strategies by the players. In some situations, players could have used a passive approach of *strategic stagnation*, wherein both players refrained from adjusting their representation of the timing intervals. The lack of strategic adjustment hindered effective cooperation between the players, impeding their ability to synchronise actions. In other situations, the players could have relied on a *paired overcompensation* strategy with a tendency to follow similar approaches or strategies. This redundancy in adjustment of the common goal could limit the range of perspectives and strategies applied to the task, potentially hindering performance. This flawed dynamic of maladaptive adjustment disrupted the delicate balance required for successful joint action. However, it is crucial to note here, that these speculative conclusions are our interpretations of the data and caution should be exercised when drawing conclusive remarks. The inter-mixed design of our paradigm poses limitations when exploring adjustment on a behavioural level. Optimising this paradigm to a block design as well as including a solo (for example with a computer) or no feedback condition could provide further support for our findings.

Cavanagh et al., (2010) observed post-error slowing after negative feedback and speed up following positive feedback of the same stimulus type (after some delay) and not immediately following feedback. They interpreted this effect as indicative of working memory for the specific stimulus type – this is not what we see here, we see a local effect of adjustment reflected in theta power modulations in the players on trials immediately after the feedback. In contrast, we did not observe the same effect for the same condition type stimuli (i.e., the global effect; refer to supplementary material “*Effects of brain-to-brain coupling on global cooperative outcomes analysis”* for further details). The difference in findings potentially lies in the methodologies: Cavanagh et al. (2010) used a probabilistic learning task in *individual* participants, whereas our participants had to take into account the behavioural adjustment of their partner as well. Post-error slowing or speeding is not applicable in the current study. Our paradigm with its random distribution of time estimation intervals means the feedback induced activity we observe is likely not reflecting lower order motor preparation processes (i.e. slowing or increasing reaction times) but rather an adjustment in their representation of the time interval. This is evident in the current results as we observed a lack of theta modulation differences between trials that necessitated a speed up vs. slow down strategy on the next trial.

Arriving at a shared representation of time through trial-and-error in a joint task can be related to Nash’s equilibrium in the context of game theory. Nash’s equilibrium is a concept that describes a state in a game where each player’s strategy is optimal given the strategies chosen by other players (Jin et al., 2012). It represents a stable outcome where no player has an incentive to unilaterally change their strategy. In the case of arriving at a shared representation of time, the participants in the joint task are engaged in a cooperative endeavour where they need to coordinate their actions based on feedback. Through trial-and-error, they aim to converge on a mutually agreed-upon understanding of ’short’, ’medium’, and ’long’ time intervals. This process involves continuous adjustments and learning from the feedback received. The relation to Nash’s equilibrium arises when the participants reach a point where their strategies, in this case, their representations of time intervals, align and are mutually optimal. It signifies that they have found a shared understanding or agreement that maximizes their joint cooperation. This equilibrium state represents a stable and satisfactory outcome in terms of their performance in the joint task.

In the context of our findings, the anti-correlated brain-to-brain activity between individuals suggests a complementary processing style, indicating that each individual is adopting different strategies or approaches. This implies that they are not in a Nash equilibrium because one individual’s adjustment in response to feedback is influencing the other individual’s subsequent performance. Nash’s equilibrium typically involves players making strategic choices based on their beliefs about the other player’s actions and seeking to maximize their own outcomes. In our study, we observed that an individual players’ brain activity and subsequent performance are influenced by the feedback received, rather than being solely driven by strategic decision-making based on beliefs about the other player’s behaviour.

As the two-brain science field continues to evolve and researchers increasingly implement neuroscientific hyperscanning methods to study social interaction, it would be important to extend the current findings pertaining the application of feedback in other social scenarios, particularly those that involve verbal communication. By doing so, we could discover valuable insights that can be applied to and improve real-world social interactions such as romantic or friendship relations, team collaborations in workplace settings or even teacher-student interactions in educational settings (Dikker et al., 2017). On the other hand, we can also delve further into the specific individual characteristics that are associated with the underlying neural processes involved in successful social interactions. For example, as raised by Sebanz et al. (2006) what is the extent to which joint action relies on Theory of Mind (ToM). Building upon our previous work demonstrating a connection between Theory of Mind and cooperation (Markiewicz et al., 2023), the next logical step is to explore the relationship between Theory of Mind abilities and the inter-person neural correlates of joint action. Furthermore, exploring individual differences in relation to the strategies employed by players and assessing the directionality of the interaction (i.e., leader versus follower) would be an intriguing avenue for future research (e.g., Konvalinka et al., 2010).

In conclusion, we studied how feedback related changes in the EEG signal between two participants involved in a non-verbal cooperative task adjusted representations of the common goal to ensure optimal joint action outcomes. We showed that following external feedback, an anti-correlated brain-to-brain activity in theta power was associated with *flexible strategic agility* and adaptive adjustment, leading to successful joint action on subsequent trials. Conversely, a correlated brain-to-brain activity in theta power of the players was linked to either a *strategic stagnation* or *paired overcompensation* strategy; both of which led to maladaptive adjustment and thus failed future joint action. By focusing on the (anti-) correlation of neural responses and moving beyond an individual-level perspective, we offer novel and more comprehensive insights into how brain-to-brain coupling predicts joint action outcomes.

## Materials and Methods

### Participants

Sixty young adults (i.e., 30 participant pairs) took part in the study. These were University of Birmingham (aged 19 to 31, *M*: 21.5, *SD*: 2.3; 12 males) students who were compensated for their time with cash payments. One participant was excluded from all analyses due to excessive EEG artefacts in the recordings; while their cooperative partner’s data were excluded from the *paired* analysis, they were kept in for the *individual* analysis.

Subsequently, we included 59 participants in the *individual* analyses, and 29 pairs for the *paired* analyses.

All participants were native English, monolingual speakers, right-handed, with normal-to-corrected vision, and no neurological or language impairments. All participant pairs reported to not know their cooperative partner in the experiment. Participants were compensated with cash for their time. Participants signed informed consent, which followed the guidelines of the British Psychology Society code of ethics and the study was approved by the Science, Technology, Engineering, and Mathematics (STEM) Ethical Review Committee for the University of Birmingham (Ethics Approval Number: ERN_19_1661).

### Cooperation task

Cooperation was measured via a non-verbal task, completed in pairs. Participants were instructed to synchronise button presses with their partner within the pair. One of three types of auditory stimuli was played simultaneously to both participants at the start of each trial using headphones (Sennheiser 289 HD). These stimuli indicated the duration of time (referred to as ‘short’, ‘medium’, or ‘long’) that participants had to wait before pressing their button and attempting to synchronise it with the other participant in the pair. Neither of the participants within a pair knew the representation of ‘short’, ‘medium’, or ‘long’ the other participant had. There was no verbal or non-verbal communication between participants to agree on a mutual representation of these time intervals - they adjusted their responses solely based on the feedback they received at the end of each trial.

Cooperation was considered successful when both participants pressed their buttons within a 250ms timeframe. On the other hand, a trial was classified as failed cooperation if the time elapsed between the two button presses was <250ms. Participants were provided with feedback regarding their performance, as explained below. We selected this timeframe threshold after conducting piloting of the paradigm (using a different set of participants) to attain an accuracy rate of approximately 50% across conditions. The purpose of setting the task goal with this threshold was to maintain an approximately equal number of trials per condition, including successful and failed joint action. Further, we included the three types of auditory cues for the same purpose. The inclusion of only one auditory cue type would make the task too trivial for participants resulting in uneven trial numbers (or even the lack of failed cooperation).

An analysis of the accuracy rates in the present dataset revealed that we succeeded in our purpose: the overall accuracy across all pairs was 56.39% (SD = 13.94). Moreover, the accuracy rates revealed that the task worked as expected, with accuracy being lower to more difficult the condition was: A Repeated measures ANOVA (Greenhouse Geisser adjusted) revealed that there was a significant effect of condition type (high/short, medium, and low/long) on accuracy (%) (*F*(1.66, 46.469) = 76.638, *p* < .001). Post-hoc Least Significant Difference (LSD) pairwise comparisons showed that the accuracy rates were significantly different for each condition (high vs. low, *p <.001*; high vs. medium, *p < .001*; medium vs. low, *p = .037*), with the high/ short condition having the highest accuracy (M = 74.586 , SD = 14.151), followed by the medium condition (M = 49.379, SD = 18.108), and with the low/ long condition having the smallest accuracy (M = 45.207, SD = 15.697).

The experiment was set up using Python 3.6 and incorporated in-house built scripts and PsychoPy functions. The experiment scripts can be downloaded from the following link: https://osf.io/ct8jb/. Both participants viewed identical instructions, stimuli, and trial presentations on separate screens with a resolution of 1920 x 1080. The button presses were recorded using Razer gaming keypad. This gaming pad uses high-precision mechanical keyboard switches and specialized internal circuitry that purportedly enables the keyboard state to be polled at a rate of 1000 times per second.

Each trial commenced with the appearance of a fixation cross, which remained on the screen for the entire duration of the trial. After 300ms, an auditory cue was played for 200ms to both participants. There were three distinct auditory cues used: (1) a high-frequency ’beep’ (2500 Hz), (2) a medium-frequency ’beep’ (1000 Hz), and (3) a low-frequency ’beep’ (200 Hz).

Participants were instructed to wait for a ’short’ amount of time upon hearing the high tone, a ’medium’ amount of time upon hearing the medium tone, and a ’long’ amount of time upon hearing the low tone. Following the auditory cue, participants were required to press their buttons and synchronize their actions. Once both participants had made their button presses, the fixation cross remained on the screen for an additional 1500ms, after which a feedback symbol was displayed. Each trial had two potential outcomes: (1) successful cooperation/joint action, indicated when the elapsed time between button presses of participants within a pair was ≤250ms, or (2) failed cooperation/joint action, indicated when the elapsed time between button presses of participants within a pair was >250ms.

For successful cooperation, both participants were shown a green smiley face as visual feedback, while failed cooperation was represented by a red sad face for both participants. This visual feedback remained on the screen for 1000ms, followed by the presentation of a fixation cross for 1000ms. Subsequently, a feedback text was displayed to both participants for 5000ms. The text varied depending on the outcome of the trial, with participants receiving either a ’Well done!’ message if their button presses were synchronized, or feedback indicating which participant pressed the button first/second and the time between the button presses (see Figure 1C).

An example of a trial presentation is depicted in Figure 1C. The experimental task consisted of 300 trials in total, divided into 10 blocks (30 trials in each block). Each block consisted of an equal number of the different auditory cues which were presented in a random (i.e., inter-mixed design) order (10 trials with a high beep, 10 trials with a medium beep, and 10 trials with a low beep). In total, the experiment included 100 trials of each auditory cue.

Participants were given the opportunity to have a break after each block.

### Procedure

Each experimental session started with the 32-electrode EEG set up of each participant. Once both participants were capped, they sat approx. 70cm from a monitor (opposite each other with a table divider in between them so that they could not see each other) (see Figure 1A for set up). Participants wore earplugs (this was to prevent hearing each other’s button presses) as well as noise cancelling headphones (to hear the auditory cues). Participants read the instructions of the task and once both participants were happy to continue, the experimenter started the ‘training’ phase.

The training phase involved participants hearing each of the auditory tones (high, medium and low) to familiarise themselves with the tones. While each tone was played, the associated description (i.e. high, medium, low) of the tone was displayed on the screen. This was repeated twice to ensure that participants were able to distinguish between the tones.

The experimenter was not present in the room during the experiment, instead the experimenter observed the participants through a one-way mirror. The experimenter entered the room upon the completion of each block and ensured that both participants were happy to continue. The experimenter controlled the start of the next block after each break.

### EEG recording

The EEG set up was the same for both participants within a pair. EEG was recorded using Waveguard caps containing 32 Ag/AgCI electrodes (10-20 layout). The EEG signal was acquired with online reference to the CPz channel. The ANTneuro EEGosports amplifier system was used to amplify the signal and the EEGo sports was used to record it. The recorded signal was sampled at 500Hz, with a 150 Hz low pass filter and a 0.05Hz high-pass filter. We aimed to keep the impedances below 10kΩ. The EEG signal of both participants was synchronized to the onset of the triggers using a parallel port splitter.

### EEG pre-processing

The EEG preprocessing was performed using EEGLAB (2019.0) (Delorme & Makeig, 2004) and Fieldtrip toolbox (2021-10-16) (Oostenveld et al., 2011). The preprocessing steps for both participants in each pair were identical. Firstly, the first 5000 data points were removed from the continuous EEG signal to remove the amplifier start up noise. The continuous signal was then filtered using a high pass filter of 0.1Hz. The notch filter was then used to eliminate the electrical noise as well as its harmonics (45 to 55Hz; 95 to 105Hz; and 145 to 155Hz).

The continuous data were visually inspected, and any noisy channels were removed and then interpolated (M = .27, SD = .67, max = 2). The signal was then re-referenced to the average of all electrodes and the CPz electrode (the online reference channel) was then added back into the data.

The data were then epoched to the onset of the feedback face (-7 to 7sec) (fourth screen in Figure 1C). Ocular artefacts were removed based on the scalp distribution using independent component analysis (ICA) in EEGLAB (2019.0). To improve the signal-to-noise ratio and classification accuracy of components, the EEG signal was high pass filtered at .8Hz (Winkler et al., 2015) for nine recordings prior to conducting ICA. ICA weights obtained from the filtered data were then added back to the original signal (filtered at 0.1 Hz). The average number of removed components was 2.88 (SD = 1.31) for each participant. For the subsequent analysis, the data were further epoched to the onset of the feedback text (the last screen of Figure 1C) (-4 to 4.8sec) and trials were sorted into meaningful conditions depending on the type of analysis carried out (see below).

## EEG analysis

### Feedback related analysis

To compare neural activity related to feedback after successful and unsuccessful cooperation, we conducted Time-Frequency representations (TFRs) of power and Event-Related Potentials (ERPs) analyses across all 59 participants. The successful condition consisted of trials (mean N = 155.64, SD = 41.08) in which both players within a pair pressed their buttons within 250ms of each other. The unsuccessful cooperation condition consisted of trials (M = 120.89, SD = 40.21) in which the time elapsed between the button presses of the players was greater than 250ms.

### Time-Frequency representations of power analysis

The TFRs of power were calculated using the ‘mtmconvol’ method in Fieldtrip. In line with previous work (Markiewicz et al., 2021) we used sliding Hanning tapers with an adaptive time window of 3 cycles per each frequency of interest. The frequency of interest range was 2-50Hz in steps of 1Hz, and the time of interest of -1 to 4.8sec in steps of .05sec locked to the onset of the feedback text of each trial. The TFRs were calculated in relation to the relative change in power from baseline, with the baseline period being -0.6sec to -0.1sec (locked to the onset of the feedback text).

The statistical significance of the differences between conditions (successful vs. unsuccessful cooperation) in feedback cue locked oscillatory power changes, specifically locked to feedback cues in the time-frequency domain, was evaluated using non-parametric cluster-based permutation tests (Maris & Oostenveld, 2007). Each channel/time/frequency data point pair (successful and unsuccessful cooperation) locked to the onset of the feedback text was compared using a dependent samples T-test with a threshold of a 5% significance level. In cases where these comparisons (successful vs. unsuccessful cooperation) exceeded the significance level, the significant pairs were then clustered based on spatial proximity (via the triangulation method – a minimum of two neighbouring electrodes were considered a cluster). The Monte Carlo simulation was then performed to obtain probability values for the clusters by randomly shuffling the condition labels 1000 times and calculating the maximum cluster level test statistic for each permutation. The permutation tests were carried out on the following pre-defined and averaged frequency range bands: delta (1-4Hz), theta (4-7Hz), alpha (8-14Hz), low beta (15-20Hz), and high beta (20-25Hz), using a time window of 0 to 4 seconds locked to the onset of the feedback text.

### Event Related Potentials analysis

The ERPs were calculated by averaging the time-locked EEG activity of all trials, separately for successful and unsuccessful cooperation conditions for each participant. The baseline correction used for the ERP analysis was -0.1sec to 0 prior to the onset of the feedback text. We employed the non-parametric cluster-based permutation tests (see above for full explanation; for comparison of ERPs the frequency part is not applicable) to statistically examine condition differences (successful vs. unsuccessful) in ERP amplitudes in a 0 to 4sec time window, locked to the onset of the feedback text. See supplementary material and Suppl. Fig. 1 for ERP results.

### Forecasting cooperative outcome analysis

#### Effects of brain-to-brain coupling on local cooperative outcomes analysis

To identify power brain-to-brain signatures between participant pairs that were predictive of *local* cooperative outcomes, we focused our analysis of the TFRs of power locked to feedback in failed joint action trials. Importantly, the TFRs analysis was performed on the same set of trials for both player one and player two within each pair. The unsuccessful trials were categorized into two conditions: *forecasting unsuccessful cooperation* on the next consecutive trial (i.e., failed cooperation trials followed by unsuccessful cooperation) and *forecasting successful cooperation* (i.e., failed cooperation trials followed by successful cooperation on the next consecutive trial). The average number of trials included in the analysis was 46.24 (SD = 14.11) for the condition of forecasting successful cooperation and 48.90 (SD = 26.95) for the condition of forecasting failed cooperation.

To compare the between player coupling of power between the two conditions, we conducted a Spearman’s Rho analysis on the power values between player one and player two for every time, frequency, and channel data points within each trial. To account for trial number inconsistencies between the conditions for each pair of participants, the Spearman rho values were converted to Fisher Z scores (Mazaheri et al., 2018). We used non-parametric cluster-based permutation tests to statistically examine condition differences (i.e., *forecasting unsuccessful cooperation* and *forecasting successful cooperation*) in the inter-player coupling of power. This is the same method as described above (in the *Time-Frequency representations of power analysis section)*, but, instead of comparing power values, we compare the Fisher Z converted correlation values (between players) at every channel/time/frequency point. These values are the “inter-player power coupling values”.

We primarily focused our permutation test analyses on the theta (4-7 Hz) band. The reasoning behind this is that theta band activity has been extensively implicated in network changes underlying behavioural adjustment and signalling the need to change strategy following an error, although this has been studied in individual brains (Cavanagh et al., 2013; Cavanagh & Frank, 2014; van de Vijver et al., 2011). We have also *exploratively* analysed the following pre-defined and averaged frequency range bands: delta (1-4Hz), alpha (8-14Hz), low beta (15-20Hz), and high beta (20-25Hz). Based on the *feedback related* results in the Time-Frequency domain, we predefined the following time windows of interest that we used in the permutation tests: 0-0.4sec and 0.6-1.05sec in the theta band, and 0.4-1.65sec in the alpha band. In addition, we exploratively used 0-2sec time windows in the permutation tests. The permutation tests identified electrode clusters that showed the greatest inter-player power coupling differences between conditions *(forecasting unsuccessful cooperation* vs. *forecasting successful cooperation)*. We also computed similar analysis to identify brain-to-brain dynamics predictive of *global* cooperative outcomes (i.e., cooperative outcomes of subsequent trials with the same condition type; see supplementary material for further details on the analysis method and the results).

### Trial-by-trial analysis linking theta dynamics with behavioural outcomes

In addition to categorizing cooperative outcomes in a dichotomous way (by using a pre-defined threshold, categorizing cooperative success within 250ms and failure beyond it), we undertook an analysis that treated the cooperative outcome as a continuous variable. The cooperative outcome as a continuous variable reflected better cooperation with a smaller absolute difference between the RT’s of the button presses of player one and two. This allowed us to examine whether the trial-by-trial inter-player theta dynamics are predictive of cooperative outcomes, when the pre-defined categories of success and failure are not present. To do this, we carried out a multiple linear regression model (estimated using Ordinary Least Squares), with the outcome variable being the *continuous* cooperative outcome (i.e., time difference between the button presses of player 1 and player 2). The predictors were the averaged feedback related individual theta power values for player one and player two. The theta power values were averaged over the time window and channels (see *Feedback related results section)*, which showed significant condition differences (successful vs. unsuccessful cooperation; 0 to 0.4sec post-feedback and over central and centro-parietal electrodes) in individual feedback cue locked oscillatory power changes. Our approach here was guided by a focus on the interaction of *individual* theta powers within a regression model rather than their pairwise correlations. Thus, we utilised the cluster of electrodes identified through the cluster-based permutation tests on feedback related oscillatory modulations at an *individual* level. The proposed model was as follows: cooperative outcome ∼ averaged theta power of player one * averaged theta power of player two + absolute theta power difference between players. This was carried out on a trial-by-trial basis. The merit of trial-by-trial analysis is that it can reveal information that would be lost if data were collapsed into a dichotomous mean of a condition (Pernet et al., 2011).

### Individual strategy adjustment analysis

To further explore the underlying mechanism responsible for the observed brain-to-brain activity in theta power forecasting joint action outcome on a local level, we conducted additional analyses. Our aim was to determine whether this effect truly originates from the *mutual* adjustment of time interval representation in joint action context, or whether it is attributed to a singular process performed independently by each individual. The rationale behind this analysis stemmed from the hypothesis that if theta band activity on *individual* level, is associated with integrating feedback into one’s actions, and considering the varying degrees of adjustments participants need to make based on their *individual* situation, correlations and anti-correlations in theta activity among individuals may arise simply due to individuals adapting their behaviour according to their own feedback, rather than any awareness of their counterparts’ actions. To address this, we categorized trials based on the type of required adjustment for the subsequent trial. Specifically, using the non-parametric cluster based permutation tests (with a pre-defined time window of 0 to .4sec based on the above results) we compared the theta (4-7Hz) power of Time-Frequency Representations (TFRs) between *unsuccessful cooperation trials that necessitated speeding up* in the subsequent trial vs *unsuccessful trials that required slowing down* in the subsequent trial. The TFRs were locked to the onset of the feedback text and collapsed across all participants (N=59).

## Data availability

Stimuli and data are available here: https://osf.io/ct8jb/

## Authorship Contributions

RM, KS and AM conceptualised the study. RM programmed the task, collected the data and analysed the data. AM and KS supervised data analyses. RM wrote the manuscript. All authors edited the manuscript.

## Acknowledgments

We would like to thank the students who helped to conduct the data collection: Rupali Limachya, Nicolas Hayston Lecaros, and Philippa Lane. We would like to thank Oscar Cass Darweish for illustrating the experimental set-up. We would also like to thank the University of Birmingham Hillary Green Studentship for funding this project.

