## Supplementary material, Supplementary Figure 1 for "Brain-to-brain coupling forecasts future joint action outcomes"

***Interplay of behavioural adjustment (between players) was predictive of cooperative outcome***

We sought to examine whether the outcome of each trial could be predicted based on the interplay of adjustments made by both players on a behavioural level. However, the nature of our experimental design, which involved random interleaving of condition types within blocks rather than a block-based structure, prevented us from directly examining sequential behavioural adjustments between consecutive trials. This constraint arose because the specific condition type consistently influenced the 'change' or 'adjustment' in reaction time (RT) due to the varying wait times required (short, medium, or long).

To overcome this, we established a reference point, or 'baseline,' to facilitate the calculation of behavioural adjustments for each trial. Our reasoning centred on the notion that 'learning' occurred immediately after the present trial, coinciding with participants receiving feedback. Consequently, for each subsequent trial following this feedback, we identified the corresponding condition type (signifying the moment of adjustment). We then located the nearest preceding trial with the same condition type, which served as our baseline. Using this approach, we computed the behavioural adjustment for each player. This adjustment was quantified as the difference between the RT of the previous trial with the same condition type and the RT of the subsequent trial with the same condition type.

With this, we were then able to fit a multiple linear regression model (estimated using Ordinary Least Squares), with the predictors being: the behavioural adjustment of player one and behavioural adjustment of player two. The outcome variable was the cooperative outcome (i.e., the time difference between the button press of player one and two); cooperative outcome ~ behavioural adjustment of player one * behavioural adjustment of player two. This was carried out on a trial-by-trial basis.

The model explained a statistically significant but weak proportion of variance (F(3, 2647) = 61.40, p < .001).The model revealed that the behavioural adjustment of player one (β = -0.14, p < .001) but not of player two (β = -1.11, p = 0.945) was a significant predictor of cooperative outcome. Most importantly, the interaction between behavioural adjustment of player one and behavioural adjustment of player two significantly predicted cooperative outcome (β = -0.16, p < .001).

Given that the labels Player 1 and Player 2 are arbitrary, we decided to focus on the interaction term, which captures the essential dynamic interplay between both players. The interaction term in a regression model captures the combined effect of two variables on the outcome. Here the significant interaction term between Player 1 and Player 2's behavioral adjustments indicates that the cooperative outcome is influenced by how both players' adjustments interact, rather than by either player's adjustment alone. This highlights the importance of mutual adjustment in successful cooperation. While this model indeed confirms the significance of the interplay between the behavioural adjustments made by both players as a strong predictor of continuous cooperative outcomes (i.e., the time gap between player one's and player two's button presses), it is important to exercise caution when interpreting these findings. Our experimental design posed limitations regarding the direct calculation of behavioural adjustments occurring from one trial to the next due to the presence of intermixed trial types, making trial type (short, medium, and long) a confounding variable. Consequently, we had to establish a baseline for behavioural adjustment using the previous trial of the same condition and calculate *behavioural adjustment*, which may not fully represent the way that participants updated their mental representations of the timing intervals on the next consecutive trial. Therefore, it is important that these results are replicated using a blocked experimental design, enabling to directly examine the effect of behavioural adjustment of both players on cooperative outcome.

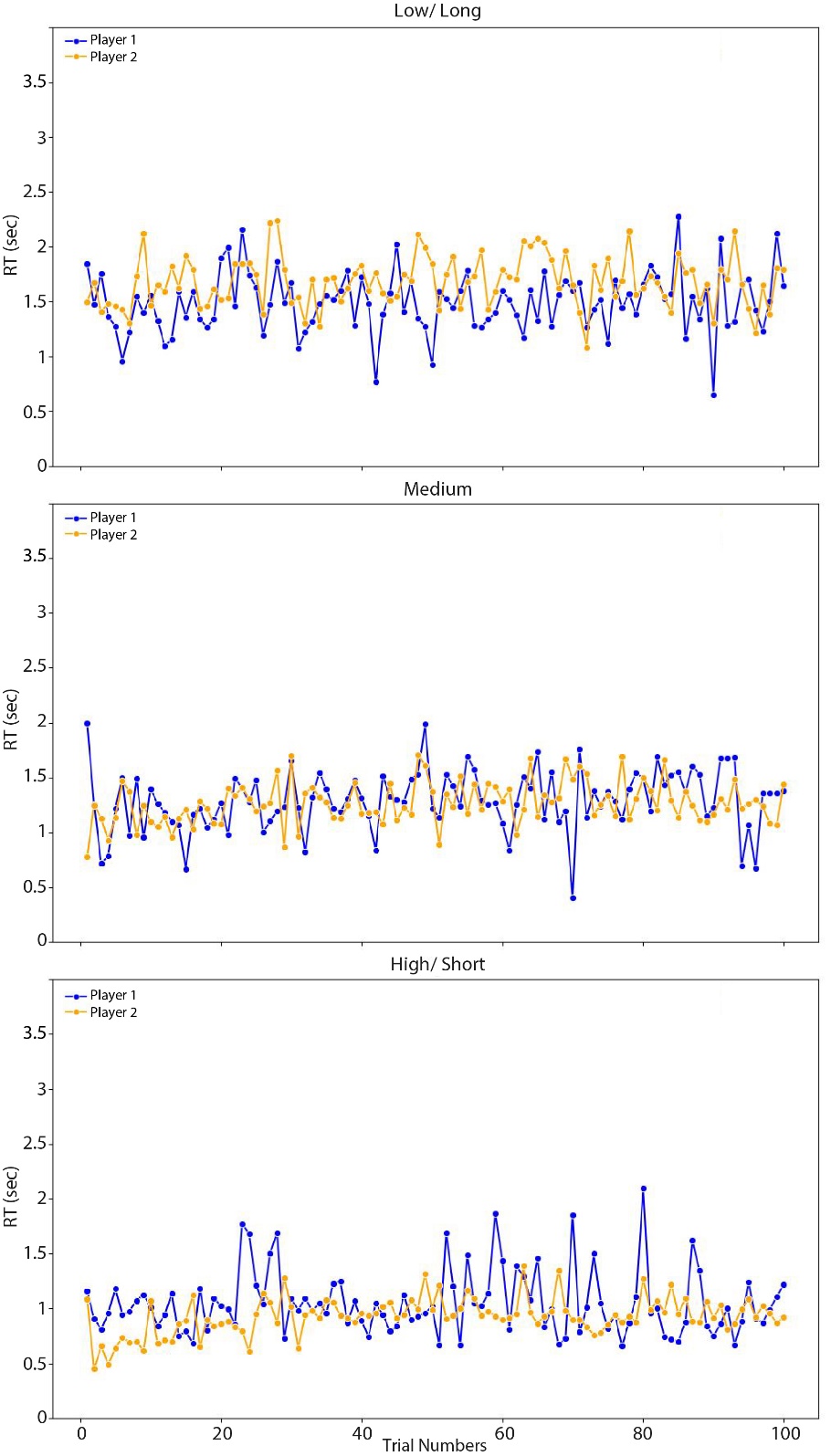

*Supplementary Figure 1.* Response times of player 1 (blue) and player 2 (orange) for pair one as an example in function of trial number (note that the trial numbers are dummy coded due to the experimental design, wherein the conditions were interleaved and randomised for each pair, and thus these represent the duration of the experiment rather than the real trial number) for each condition (*top row*: low, *middle row*: medium, and *bottom row*: high). A comprehensive set of similar figures (for other pairs) can be found on: https://osf.io/ct8jb/. We also provide the code used for plotting along with dataset itself, enabling users to adjust the axis limits as needed, particularly in case of outlying responses.

***Participants were more likely to successfully cooperate if the previous trial (of the same type) was also successful***

At the behavioural level, we investigated whether pairs are more likely to maintain successful cooperation if the previous trial of the same type was also successful. We used a 2 (trial type: Correct followed by correct vs. incorrect followed by correct) x 3 (condition: short, medium, and long) repeated measures ANOVA to compare the frequency of trials where a successful cooperation was followed by another successful trial against those where an unsuccessful cooperation was followed by a successful one for each condition type (short, medium, and long). Suppl. Fig. 2 displays the average frequency of successful and unsuccessful trials followed by successful trials for each condition type. We found that there was a significant main effect of trial type, F(1, 28) = 40.23, p < .001, revealing that there was a greater frequency of correct followed by correct trials (M=37.5, SD=23.1) compared to incorrect followed by correct trials (M=18.6, SD=5.27). There was also a significant main effect of condition F(1.65, 46.20) = 76.26, p < .001, and a post hoc one-way ANOVA showed that pairs were more likely to successfully cooperate following a previous successful trial compared to when the previous trial was unsuccessful in the short (M difference = 43.93, F(1, 28) = 99.90, p < .001) and medium (M difference = 9.76, F(1, 28) = 7.64, p = .010) conditions, but this was not observed in the long condition (M difference = 2.93, F(1, 28) = 1.05, p = .314). There was also a significant interaction between trial type and condition, F(1.58, 44.23) = 71.91, p < .001. The results from pairwise comparisons are displayed in Suppl. Table 1.

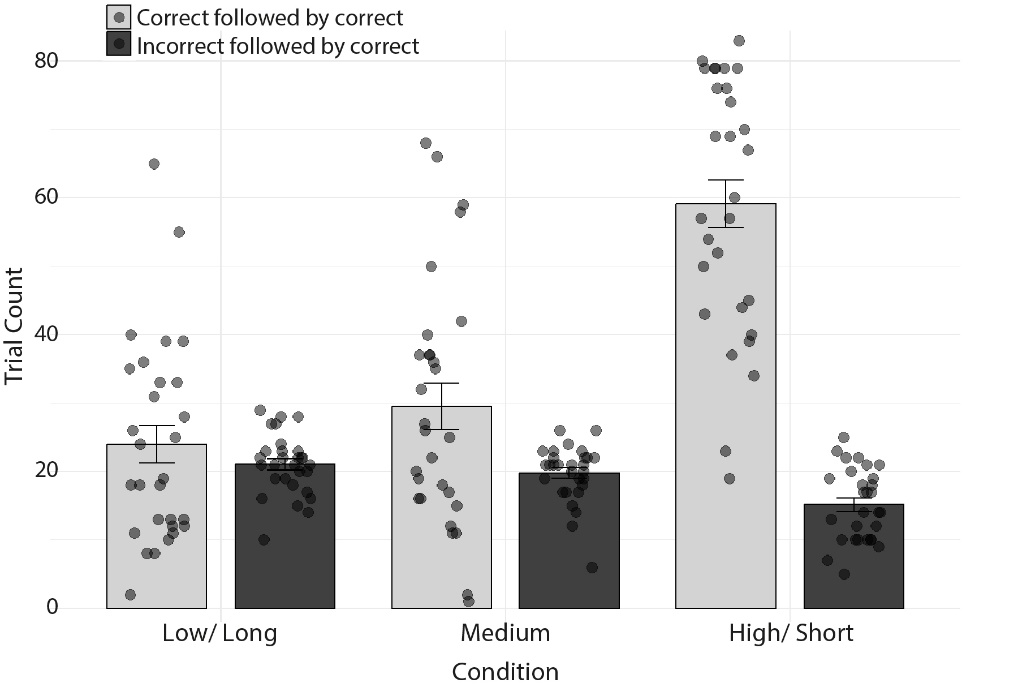

*Supplementary Figure 2.* The average frequency of trials where successful cooperation is followed by another successful cooperation (light grey) compared to trials where unsuccessful cooperation is followed by successful cooperation (dark grey) across each condition type (long, medium, and short). Individual data points are represented by black dots, and error bars indicate the standard error.

Supplementary Table 1. *Pairwise t-test comparisons for the frequency of trials where a successful cooperation was followed by another successful trial (Correct followed by Correct) against those where an unsuccessful cooperation was followed by a successful trial (Incorrect followed by Correct) between conditions (short, medium, and long). Adjusted p indicates the p-value after Bonferroni correction.*

| **Trial type** | **Condition(1)** | **Condition(2)** | **N(1)** | **N(2)** | **t** | **df** | **P** | **Adj. p** |
| --- | --- | --- | --- | --- | --- | --- | --- | --- |
| Correct followed by Correct | Long | Medium | 29 | 29 | -2.60 | 28 | .015 | .045 |
| Correct followed by Correct | Long | Short | 29 | 29 | -10.30 | 28 | <.001 | <.001 |
| Correct followed by Correct | Medium | Short | 29 | 29 | -9.20 | 28 | <.001 | <.001 |
| Incorrect followed by Correct | Long | Medium | 29 | 29 | 1.68 | 28 | .105 | .315 |
| Incorrect followed by Correct | Long | Short | 29 | 29 | 4.65 | 28 | <.001 | <.001 |
| Incorrect followed by Correct | Medium | Short | 29 | 29 | 4.13 | 28 | <.001 | <.001 |

***The trial number of first successful cooperation varied across condition types***

We examined the differences in the trial numbers of the first successful cooperation among the different conditions (long, medium, and short). A one-way ANOVA revealed a significant effect of condition on the trial number at which the first successful cooperation occurred (F(2, 84) = 3.85, p = 0.025). Suppl. Fig. 3 shows the average trial number of first successful cooperation across conditions. Post-hoc Tukey tests indicated a significant difference between the medium and short conditions, showing that the first instance of successful cooperation occurred earlier in the short condition compared to the medium condition (p = .039). There was a marginal effect when comparing the long and short conditions, with the first instance of successful cooperation tending to occur earlier in the short condition compared to the long condition (p = .059). No significant difference was found between the long and medium conditions (p = .985).

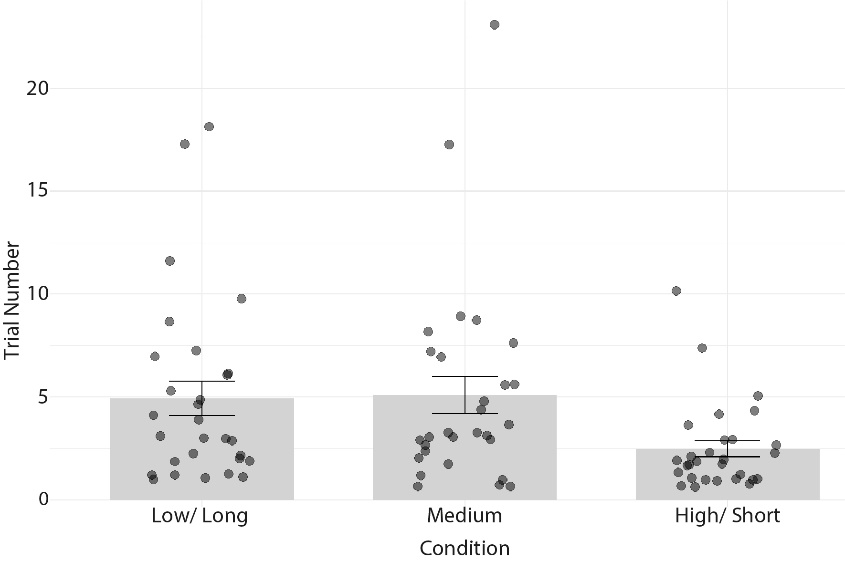

*Supplementary Figure 3.* The average trial number at which the first successful cooperation occurred for each condition type (long, medium, and short). Individual data points are represented by black dots, and error bars indicate the standard error.

***Feedback related results***

*Feedback related oscillatory modulations in delta, theta, alpha and beta bands*

We compared the feedback related activity, which includes evoked and induced activity, between correct and incorrect conditions across all of the participants (N =59) aligned to the onset of the feedback text. Suppl. Figure 4A displays the TFRs of power locked to the onset of the feedback text for the incorrect, and correct conditions, the difference between them, as well as the topographic distributions of the condition effects. Below we describe the significant oscillatory differences between correct (i.e., successful cooperation) and incorrect (i.e., failed cooperation) conditions that were identified using non-parametric cluster-based permutation tests using a 0-4sec time window post feedback text onset. We observed significant condition differences in the pre-defined delta (1-4Hz), theta (4-7Hz), alpha (8-14Hz), low beta (15-20Hz) and high beta (20-25Hz) frequency bands following the onset of the feedback text. Firstly, we observed significant condition differences (p = .003 and p <.001 respectively) in theta and delta activity maximal over the central and centro-parietal channels. Specifically, theta (and delta) power was stronger right after the onset of the feedback text in the incorrect condition until around 0.4sec compared to the correct condition. It has been previously proposed that mid-frontal theta activity serves as a neural indicator of monitoring ongoing actions and signalling unfavourable action outcomes. This in turn results in adaptive response and modulated activity due to committed errors (Beldzik et al., 2022; Mazaheri et al., 2009). Further, there was a significant condition difference (p < .001), with greater delta power in the incorrect vs. correct condition. This effect corresponded to a cluster maximal over the central electrodes that stretched between 1.15 to 1.75sec post feedback text onset. We consider this effect to be a representation of the sustained ERP in the similar time window. Finally for the delta band, we found significantly more pronounced activity in the incorrect compared to the correct condition following the feedback onset (p = .006). This effect was most prominent between 2.65sec and 2.75sec post feedback text onset and maximal over right centro-temporal channels. Further, in the theta band, there was a more sustained theta attenuation in the incorrect compared to the correct condition (p < .001). This effect was maximal over centro-parietal and parietal sites and most prominent between 0.6 to 1.15sec post feedback text onset. We take this as spectral leakage from the effect seen in the alpha band, see below for the interpretation of this effect. Lastly for the theta band, we observed a condition effect that was maximal over temporal electrodes and left lateralised (p = .039). Specifically, there was a reduction in theta power in the incorrect condition relative to the correct condition corresponding to a cluster that stretched between 1.5 and 1.6sec post feedback onset. We also found significant condition differences in alpha activity. Firstly, there was a greater alpha power decrease in the incorrect condition relative to the correct condition (p < .001). This effect corresponded to a cluster that extended between 0.3 to 1.75sec post feedback and was maximal over central electrodes. We interpret this post error alpha decrease as stronger engagement, greater resource allocation and active processing to prevent future errors. This enhanced and more alert cognitive state optimises visual processing and better motor system coordination for future trials (Mazaheri et al., 2009). We also found that there was a greater alpha power increase in the correct condition relative to the incorrect condition (p = .031). This effect corresponded to a cluster that spread between 2 to 2.25sec and was maximal over left temporal channels.

We found a significant condition effect in the low beta band (p =.024). This corresponded to a transient time window of .2 to .25sec post feedback and was maximal over centro-parietal electrodes. Further, there was a prolonged opposite pattern of low beta (p < .001) and high beta (p < .001) activity between the conditions. The incorrect condition yielded a prolonged low beta decrease, whereas the correct condition showed a prolonged low beta increase. This effect corresponded to a cluster that spanned from .35 to 1.5sec and was prominent all over the scalp. The beta power increase following correct responses and thus feedback is likely reflecting the requirement for strategy maintenance by facilitating the reinforcement of the current motor and cognitive strategy (Cohen et al., 2007; Engel & Fries, 2010; Marco-Pallares et al., 2008, 2009; Yeung et al., 2004). Lastly, we found a transient condition effect (p = .013) in the high beta band that extended between 2.9 and 3sec post feedback. Specifically, there was a greater high beta power increase in the incorrect compared to the correct condition, maximal over bilateral temporal sites.

*Initial sustained ERP positivity and later negativity associated with incorrect trials after feedback*

We compared the ERPs between incorrect and correct conditions across all of the participants (N =59) using a time window of 0 to 4sec post feedback text onset. Suppl. Figure 4B shows feedback locked averaged ERP waveforms for incorrect and correct conditions, as well as the condition effect topographic distributions. The cluster-based permutation tests revealed that there was a significant effect of condition, specifically the incorrect condition was associated with a greater positivity compared to the correct condition (p < .001). This effect corresponded to a cluster that spanned from .178 to .926sec and was maximal over centro-parietal electrodes. Secondly, we found another condition effect, where the incorrect trials yielded a sustained negativity ERP relative to the correct trials (p < .001, p = .034). This corresponded to clusters that extended from 1.406 to 2.554sec as well as 2.562 to 2.634sec respectively, most prominent over centro-parietal channels. Previous research has proposed that the localization of error-related negativity (ERN) and theta activity overlap, suggesting a shared involvement in error monitoring processes (Mazaheri et al., 2009). Additionally, it has been suggested that the ERN serves not only as an error detection mechanism but also predicts the extent of learning from errors (Frank et al., 2005). Consequently, we interpret this ERP effect playing a role in error monitoring and adjustment for future actions.

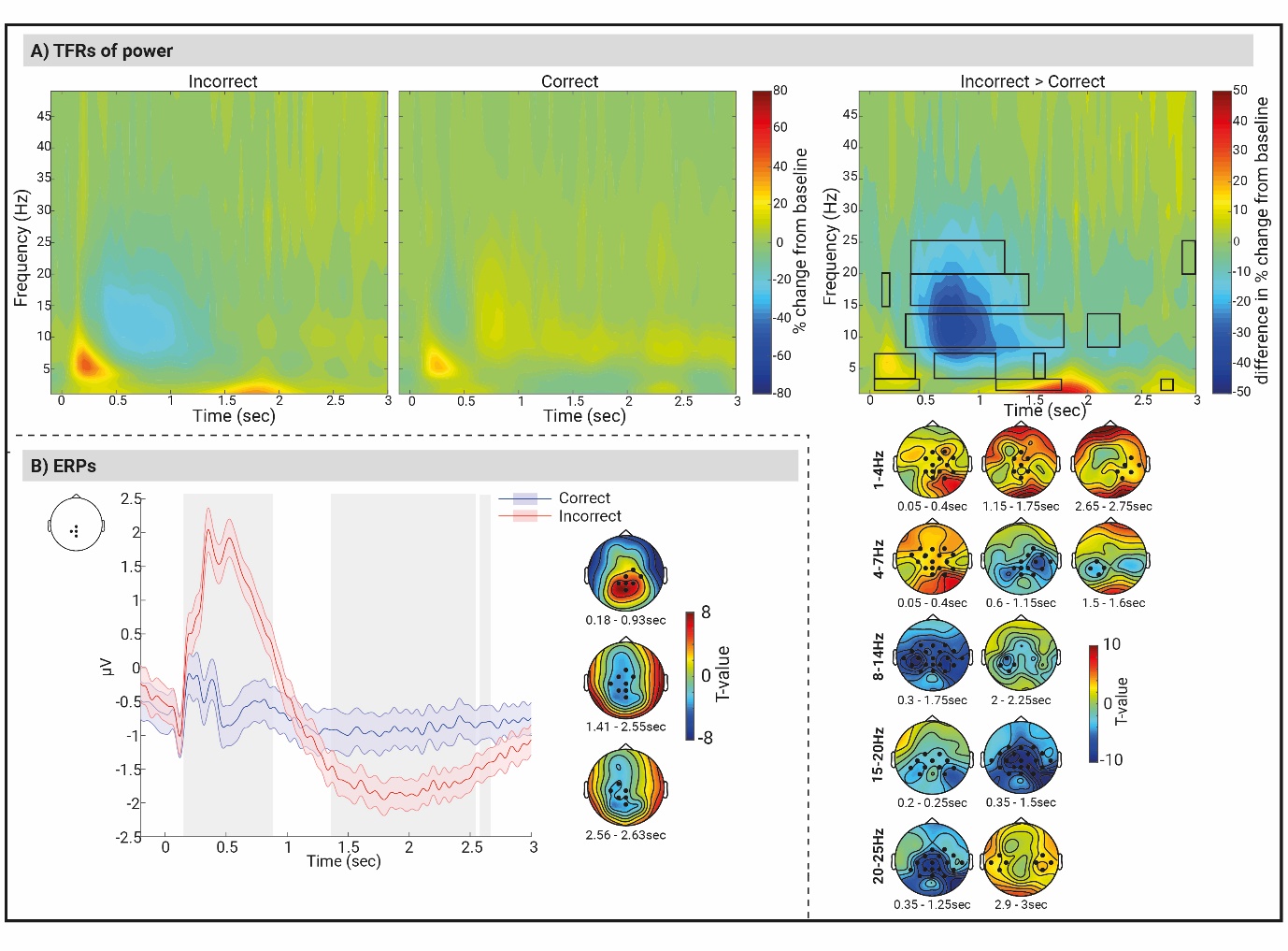

*Supplementary Figure 4.* (A) TFRs of power (at the Cz electrode) for incorrect trials (left), correct trials (middle), and the condition differences (i.e., incorrect – correct) (right) averaged over all individuals (N = 59), locked to the onset of feedback text. Headplots illustrate the clusters of electrodes that show the most pronounced mean condition difference. Black rectangles indicate significant condition differences (p < 0.05, cluster corrected). (B) Feedback locked averaged ERPs produced by correct (blue) and incorrect (red) trials in the timing task averaged over all individuals (N = 59). The ERP waveforms show averaged ERPs across the overlapping (across the three time windows of the significant between condition effects) electrode clusters (Cz, CP1, Pz, and CPz) that indicate the maximal condition difference (a schematic view of these electrodes is shown in the top right corner). The shaded areas around the ERP waves represent standard error. The grey rectangles represent time windows of the significant between condition differences. The black dots in the headplots illustrate the clusters of electrodes showing most pronounced mean condition differences for each time window.

***Effects of brain-to-brain coupling on global cooperative outcomes analysis***

We performed further analysis to identify power brain-to-brain signatures between participant pairs that were predictive of *global* cooperative outcomes. In our experimental design, condition types (i.e., high/short, medium, and low/long) were interleaved in a randomised manner within each experimental block, meaning that successive trials frequently did not share the same condition type. Given this setup, we aimed to explore the potential for predicting cooperative outcomes of subsequent trials with the same condition types based on inter-player neural signatures – we refer to as the "global effect".

Our approach focused on examining the nearest trial of the same condition type, instead of studying consecutive trials. Specifically, we categorized unsuccessful trials into two distinct conditions: forecasting unsuccessful cooperation on the nearest trial of the same condition type (involving failed cooperation trials followed by another unsuccessful cooperation trial of the same condition type), and forecasting successful cooperation on the nearest trial of the same condition type (entailing failed cooperation trials followed by a successful cooperation trial in the subsequent non-consecutive instance). The average number of trials included in the analysis was 41.28 (SD = 12.27) for the condition of forecasting successful cooperation and 53.45 (SD = 29.58) for the condition of forecasting failed cooperation. In order to compare the two conditions we used non-parametric cluster-based permutation tests across all pairs (N = 29 pairs). We followed the exact same steps as described in the main manuscript section “*Effects of brain-to-brain coupling on local cooperative outcomes analysis”*.

*Inter-player power dynamics did not forecast cooperative results in the following trial of the same condition*

No significant inter-player power coupling patterns were observed when forecasting cooperative outcomes for the nearest trial of the same condition type within the theta band (utilizing the predefined time window of 0-0.4 seconds), nor within the alpha band (utilizing the predetermined time window of 0.3 to 1.75 seconds). Subsequent exploratory analyses involving a time window of 0 to 2 seconds post feedback similarly failed to reveal significant effects across any of the examined frequency bands, including alpha, low beta, and high beta.

Beldzik, E., Ullsperger, M., Domagalik, A., & Marek, T. (2022). Conflict- and error-related theta activities are coupled to BOLD signals in different brain regions. *NeuroImage*, *256*(February), 119264. https://doi.org/10.1016/j.neuroimage.2022.119264

Cohen, M. X., Elger, C. E., & Ranganath, C. (2007). Reward expectation modulates feedback-related negativity and EEG spectra. *NeuroImage*, *35*(2), 968–978. https://doi.org/10.1016/j.neuroimage.2006.11.056

Engel, A. K., & Fries, P. (2010). Beta-band oscillations-signalling the status quo? *Current Opinion in Neurobiology*, *20*(2), 156–165. https://doi.org/10.1016/j.conb.2010.02.015

Frank, M. J., Woroch, B. S., & Curran, T. (2005). Error-related negativity predicts reinforcement learning and conflict biases. *Neuron*, *47*(4), 495–501. https://doi.org/10.1016/j.neuron.2005.06.020

Marco-Pallares, J., Cucurell, D., Cunillera, T., Garcia, R., Andres-Pueyo, A., Munte, T. F., & Rodriguez-Fornells, A. (2008). Human oscillatory activity associated to reward processing in a gambling task. *Neuropsychologia*, *46*(1), 241–248. https://doi.org/10.1016/j.neuropsychologia.2007.07.016

Marco-Pallares, J., Cucurell, D., Cunillera, T., Krämer, U. M., Càmara, E., Nager, W., Bauer, P., Schüle, R., Schöls, L., Münte, T. F., & Rodriguez-Fornells, A. (2009). Genetic Variability in the Dopamine System (Dopamine Receptor D4, Catechol-O-Methyltransferase) Modulates Neurophysiological Responses to Gains and Losses. *Biological Psychiatry*, *66*(2), 154–161. https://doi.org/10.1016/j.biopsych.2009.01.006

Mazaheri, A., Nieuwenhuis, I. L. C., Van Dijk, H., & Jensen, O. (2009). Prestimulus alpha and mu activity predicts failure to inhibit motor responses. *Human Brain Mapping*, *30*(6), 1791–1800. https://doi.org/10.1002/hbm.20763

Yeung, N., Botvinick, M. M., & Cohen, J. D. (2004). The neural basis of error detection: Conflict monitoring and the error-related negativity. *Psychological Review*, *111*(4), 931–959. https://doi.org/10.1037/0033-295X.111.4.931
